# DKW basal salts improve micropropagation and callogenesis compared to MS basal salts in multiple commercial cultivars of *Cannabis sativa*

**DOI:** 10.1101/2020.02.07.939181

**Authors:** Serena R.G. Page, Adrian S. Monthony, A. Maxwell P. Jones

## Abstract

Micropropagation of *Cannabis sativa* is an emerging area for germplasm storage and large-scale production of clean plants. Existing protocols use a limited number of genotypes and are often not reproducible. Previous studies reported MS + 0.5 μM TDZ to be optimal for *Cannabis* nodal micropropagation, yet our preliminary studies using nodal explants suggested this media may not be optimal. It resulted in excessive callus formation, hyperhydricity, low multiplication rates, and high mortality rates. Following an initial screen of four commonly used basal salt mixtures (MS, B5, BABI, and DKW), we determined that DKW produced the healthiest plants. In a second experiment, the multiplication rate and canopy area of explants grown on MS + 0.5 μM TDZ and DKW + 0.5 μM TDZ were compared using five drug-type cultivars to determine if the preference for DKW was genotype-dependent. Four cultivars had significantly higher multiplication rates on DKW + 0.5 μM TDZ with the combined average being 1.5x higher than explants grown on MS + 0.5 μM TDZ. The canopy area was also significantly larger on DKW + 0.5 μM TDZ for four cultivars with the combined average being twice as large as the explants grown on MS + 0.5 μM TDZ. In the third experiment, callogenesis was compared using a range of 2,4-D concentrations (0-30 μM) on both MS and DKW and similarly, callus growth was superior on DKW. This study presents the largest comparison of basal salt compositions on the micropropagation of five commercially grown *Cannabis* cultivars to date.

## Introduction

As the commercial *Cannabis* industry goes through a rapid period of growth, many growers are challenged by a lack of a uniform, true-to-type seed for most commercially desirable cultivars. Instead, commercial cultivation of *Cannabis* is dependent on vegetative propagation from mother plants to ensure a genetically consistent crop (Chandra et al. 2017). Maintaining mother plants requires considerable production space (up to 15% of total growth space) while also risking infection of the mother plants and the subsequent propagules by pests, disease, and viruses. This, combined with the limited number of pesticides registered in the *Cannabis* industry, makes clean starting material vital for commercial production. With an effective *in vitro* protocol, multiple mother plants could be maintained for years with little risk of pest or disease using significantly less space while providing a consistent supply of clean plants (George et al. 2008; Chandra et al. 2017).

Fewer legal constraints surrounding the use of industrial hemp (*Cannabis sativa* with <0.3% w/w THC in the flowering heads) (Government of Canada 2015) have meant that early micropropagation studies often relied on its use. These early studies can be traced back to the 1980s, and focused on assessing the recalcitrance of *Cannabis* to tissue culture (Richez-Dumanois et al. 1986). Due to prohibition, early studies were limited in number and scope. However, in the last decade *Cannabis* tissue culture has seen a significant increase in the number of publications addressing a variety of aspects including regeneration, synthetic seed production and screening for elite drug-type (high THC and/or high CBD) cultivars (Lata et al. 2009b, 2010; Piunno et al. 2019). Despite the increasing number of published *Cannabis* micropropagation protocols, there is only one *in vitro* study of drug-type *C. sativa* which has included multiple cultivars in its development (Piunno et al. 2019). There is also debate on whether *Cannabis* should be designated as monospecific or polyspecific due to how polymorphic the genus is (Small 1975; Small and Cronquist 1976; Hillig 2004, 2005; Clarke and Merlin 2013; McPartland 2018). Settling this debate is beyond the scope of our study, however we will adopt the monospecific framework in this study. Regardless of the approach adopted, existing literature reveals an important lack in cultivar diversity when developing micropropagation protocols. Given the highly diverse and difficult to track pedigree of the plants of the genus *Cannabis* more genetic diversity must be included in contemporary *Cannabis* studies to ensure the results are not genotype specific.

Contemporary studies of *Cannabis* tissue culture have relied almost exclusively on media made with Murashige & Skoog basal salts (MS) (Murashige and Skoog 1962; Chandra et al. 2009; Lata et al. 2010, 2016; Farag 2014; Movahedi et al. 2015; Chaohua et al. 2016; Piunno et al. 2019; Smýkalová et al. 2019). This basal medium has been used in many studies, and one of the most successful protocols found that MS supplemented with 0.5 μM thidiazuron (TDZ) was ideal for drug-type *Cannabis* (Chandra et al. 2009, 2011; Lata et al. 2009b, 2010; Movahedi et al. 2015; Chaohua et al. 2016; Piunno et al. 2019). However, no studies exist which have offered a direct comparison between many of the commonly used basal salt compositions. Preliminary studies in our lab found that despite its reported success, MS basal salts + 0.5 μM TDZ (hereafter referred to as MS-T05) produced cultures which were not as healthy and had much lower multiplication rates than reported in the literature. Plants grown on MS-T05 exhibited high levels of callusing, hyperhydricity, signs of nutrient deficiency, low multiplication rates, and ultimately led to plant death in some cultivars. In the present work, we observed the vegetative growth of five drug-type cultivars and callus development of one drug-type cultivar on media with various basal salt mixtures. We observed that vegetative plant response varied considerably among the five drug-type cultivars tested with a preference amongst most cultivars for DKW basal salts, a trend that was also reflected in callogenesis. Due to these responses, we hypothesized that MS basal salt is not ideal for the micropropagation of *Cannabis* and that an improved DKW-based medium would increase plant growth and facilitate micropropagation and callogenesis of a wider range of *Cannabis* genetics.

## Materials and Methods

### Plant Material

Five of our cultivars, BA-21, BA-41, BA-49, BA-61, and BA-71 were supplied by Canopy Growth Corporation, and one cultivar, BA-1 was supplied by HEXO Corp. The cultivars were given code names at the request of the companies for privacy reasons. These cultivars have been maintained in culture in our facility for over a year. In preparation for these experiments, the Canopy cultivars were maintained on media consisting of DKW Basal Medium with Vitamins, 3% sucrose (w/v), and 0.7% agar (w/v) for a month to prevent any lingering effects from plant growth regulators (PGRs). The HEXO cultivar was also maintained on media consisting of DKW Basal Medium with Vitamins, 3% sucrose (w/v), 0.6% agar (w/v), and 1 mL/L PPM for a month. *In vitro* grown fresh stem tissue was used for all DNA extractions. For each cultivar (BA-1, BA-21, BA-41, BA-49, BA-61, and BA-71) three, 1 cm stem segments, each obtained from three separate clonal explants, were pooled together. The approximate weight of each of these fresh tissue samples was 100 mg.

### DNA Extraction and DNA Barcoding

DNA extraction and amplification was performed using the standard protocols for plant DNA barcoding published by the Canadian Centre for DNA Barcoding (CCDB) at the University of Guelph (Kuzmina and Ivanova 2011a; Kuzmina et al. 2011). Final DNA concentration was 20-40 ng/μL and markers were amplified using Platinum^®^ *Taq* DNA polymerase (Invitrogen) as outlined in (Kuzmina and Ivanova 2011a). DNA-barcodes were constructed using the chloroplast maturase K gene (*matK*) primer sets (Kuzmina and Ivanova 2011b). Products were analyzed using an ABI3730xl capillary sequencer (Applied Biosystems) as outlined by the CCBD (Ivanova and Grainger 2007). All sequences were blasted against both NCBI and BOLD databases for species identification.

### DNA Extraction and SSR Sequencing

DNA extractions were performed using NucleoSpin^®^ Plant II Mini (Macherey-Nagel) as per manufacturer’s instructions with the following minor modifications: PL1 and RNase A buffers were added to tissue samples in a 2.0 mL microcentrifuge tube prior to homogenizing. Samples were homogenized using a SPEX SamplePrep 1600MiniG^®^ homogenizer (SPEX) at 1500 RPM for 1 minute. Following homogenization lysate was settled by centrifugation. Samples were gently resuspended and incubated for 15 minutes at 65 °C and extraction was subsequently performed as per manufacturer’s instructions. Extracted DNA was stored at −20 °C prior to SSR genotyping. Purified DNA was thawed at 4 °C and gently vortexed prior to quantification. DNA purity and concentration were measured with a NanoDrop ND-1000 spectrophotometer (Thermo Scientific). Eleven SSR markers developed by Alghanim & Almirall were used for genotyping (Alghanim and Almirall 2003). PCR amplification was based on the Schuelke method (Schuelke 2000). Each PCR reaction consisted of: 3 μL 20% trehalose, 4.92 μL of molecular-grade H_2_O, 1.8 μL 10X PCR buffer (MgCl_2_), 3 mM dNTP mix, 0.12 μL of 4 μM M13-tailed forward primer, 0.48 μL of 4 μM reverse primer, 0.48 μL of 4 μM (universal) M13 primer labeled with VIC fluorescent dye (Applied Biosystems), 0.2 μL of 5.0 U μL^−1^ *taq* polymerase (Sigma Jumpstart™, Sigma-Aldrich), and 3 L of template DNA for a total reaction volume of 15 μL. Amplification reactions were carried out using thermocyclers (Eppendorf MasterCycler^®^). The conditions of the PCR amplification are as follows: 94 °C (5 min), then 30 cycles at 94 °C (30 s)/56 °C (45 s) /72 °C (45 s), followed by 8 cycles at 94 °C (30s)/53 °C (45S)/ 72 °C (45 s), followed by a final extension at 72 °C for 10 minutes. Plates were maintained at 4 °C at the cycle’s end and subsequently stored at −20 °C until sequencing. Fragment analysis of the completed PCR products was done using an Applied Biosystems^®^ 3500 Genetic Analyzer (ThermoFisher). A dye-labeled size standard (GeneScan 500-LIZ, Life Technologies) was used as the internal size standard, and PCR fragment sizes were quantified using a DNA fragment analysis software (GeneMarker, SoftGenetics LLC).

SSR sequencing data was processed using GeneMarker (SoftGenetics LLC). Results of the SSR allele-call analysis were organized in Microsoft Excel (Microsoft Corp.) and imported into GraphicalGenoTyping (GGT 2.0) and a relative genetic distance matrix was developed to compare pair-wise distance values. This matrix was visually represented as a NJ dendrogram (Fig. S1).

### Growth Chamber Conditions

For all the described experiments and stock material, the plants were grown within a controlled environment walk-in growth chamber. The chamber’s conditions were set at 25 °C under a 16-hour photoperiod. Photosynthetically active radiation (PAR) and light spectral data were obtained using an Ocean Optics Flame Spectrometer (Ocean Optics). The average PAR of the experimental area was 41 ± 4 μmol s^−1^ m^−2^ (Fig. S2).

### Preliminary Basal Salt Screening

Four basal salts were screened: MS Basal Salt Mixture, DKW Basal Salt Mixture, BABI Basal Salt Mixture, and Gamborg’s B5 Basal Salt Mixture (all obtained from Phytotechnology Laboratories). To ensure the results were due solely to the basal salt mixture, each treatment included a basal salt, 0.7% agar (w/v) (Fisher Scientific), 3% sucrose (w/v), 1 mL/L of B5 vitamins (Phytotechnology Laboratories), and were pH adjusted to 5.7 using 1 N NaOH. For each treatment, 200 mL of media was poured into a We-V Box (We Vitro Inc.) and autoclaved for 20 minutes at 121 °C and 18 PSI.

There were five biological replicates per treatment (boxes; n=5) with five pseudoreplicates which were 2-node stem explants from the *Cannabis* cultivar BA-1 (HEXO Corp.). Two-node explants were used instead of single-node explants since our previous experience found that single-node explants have a lower survival rate. The screening used a completely randomized design with the boxes placed on the same shelf within a controlled environment growth chamber. After 30 days of growth, each box was photographed, and qualitative observations were made. The photographs were taken in a photo booth under an incandescent lightbulb with a Canon EOS 70D. Qualitative observations were made for general explant health, leaf morphology, and hyperhydricity. Explants were determined to be hyperhydric if they appeared translucent and saturated with water and had abnormal leaf morphology.

### MS and DKW Vegetative Growth Comparison

To compare the multiplication rate and canopy growth of explants grown on MS Basal Salt Mixture or DKW Basal Salt Mixture (both obtained from Phytotechnology Laboratories), both treatments contained: 0.7% agar (w/v) (Fisher Scientific), 3% sucrose (w/v), 1 mL/L of B5 vitamins (Phytotechnology Laboratories), and 0.5 μM TDZ (Fisher Scientific) and were pH adjusted to 5.7 using 1 mM NaOH. For each treatment, 200 mL of media was poured into a We-V Box (We Vitro Inc.) and was then autoclaved for 20 minutes at 121 °C and 18 PSI. B5 vitamins were added to ensure the results were due solely to the basal salt mixtures and not the differing levels of vitamins. The two treatments shall henceforth be referred to as MS-T05 and DKW-T05 respectively.

To evaluate genotypic variability regarding the effect of the basal salt mixtures, five unique *Cannabis* cultivars were used. All cultivars were provided by Canopy Growth Corporation. The cultivars used were: BA-21, BA-41, BA-49, BA-61, and BA-71 (Fig. S1) and were sampled from *in vitro* grown mother plants as described previously.

Overall, there were three separate trials for this experiment, the first trial was run with BA-21 using five biological replicates per treatment (boxes; n=5) with five pseudoreplicates (2-node explants) each. The second trial was run with BA-41 with four biological replicates per treatment (n=4) and BA-49 with six biological replicates per treatment (n=6), both cultivars had four pseudoreplicates per replicate. The third trial was run with BA-61 and BA-71 with five biological replicates per treatment (n=5) with five pseudoreplicates each. The varying replicate and pseudoreplicate numbers between trials were due to the limited availability of plant material for certain cultivars. For each trial, all stem explants had two nodes and all boxes included one apical explant with two nodes to ensure that the explant source did not affect growth rates. Each trial used a completely randomized design and the boxes were placed on the same shelf within a controlled environment growth chamber.

### Statistical Analysis

To determine the canopy area, culture vessels were photographed from above at the end of the experiment under incandescent lighting with a Canon EOS 70D camera. Pictures were processed using ImageJ 1.50i software (National Institute of Mental Health). In brief, the outer edge of the canopy of each living explant was traced using the Freehand Line tool and the canopy area was determined from this shape using the Analyze>Measure function. Scale was set by the inclusion of a ruler in each digital photograph. All statistical analyses were performed using SAS Studio software (v9.4; SAS Institute Inc.). The data was collected when plants reached the top of the vessel. Due to differing growth rates among cultivars, observations for BA-21 were taken after 43 days, observations for BA-41 and BA-49 were taken after 41 days, and observations for BA-61 and BA-71 were taken after 35 days.

The means for the biological replicates were obtained by averaging the surviving pseudoreplicates and were used for the statistical analyses. The multiplication rate was measured by subculturing each box and counting the number of two-node explants produced from the original starting explants.

### MS and DKW Callogenesis Comparison

Callusing media were prepared with four different concentrations of 2,4-dichlorophenoxyacetic acid (2,4-D; Fisher Scientific), either 0, 10, 20 or 30 μM, on each of two basal salts: MS Basal Salt Mixture and DKW Basal Salt Mixture (both from PhytoTechnology Laboratories). Media consisted of either DKW or MS salts, 3% sucrose (w/v), 1 mL/L PPM (Plant Cell Technology), 0.6% agar (w/v) (Fisher Scientific) and pH was adjusted to 5.7 using 1 mM NaOH. The media were autoclaved for 20 minutes at 121 °C and 18 PSI. Approximately 25 mL of the autoclaved media were dispensed into sterile 100 x 15 mm Petri dishes (Fisher Scientific) in a laminar flow hood (DFMZ). Leaf explants were taken from the *Cannabis* cultivar BA-1 (HEXO Corp.) and were cultured onto the Petri dishes and placed using a completely randomized design on the same shelf within a controlled environment growth chamber.

This experiment was performed as two independent trials. Three biological replicates were prepared per treatment level in the first experiment (Petri dish; n=3), while four biological replicates were used in the second trial (n=4). Each biological replicate contained six pseudoreplicates (1 x 1 cm leaf explants).

### Statistical Analysis

The development of callus on each Petri dish was photographed weekly to measure the surface area of callus produced using ImageJ 1.50i software (National Institute of Mental Health) as described previously with canopy area. Callus mass was obtained 35 days post-induction using an electronic balance (Sartorius Quintix2102-1S). The means for the biological replicates were obtained by averaging pseudoreplicates. All summary statistics and statistical analyses were performed in Prism (v8.3; GraphPad, Inc.) using the means of the biological replicates (n=3, trial 1; n=4, trial 2).

## Results

The SSR analysis shows that all tested cultivars are genetically unique with two predominant clusters forming (Fig. S1). DNA barcoding of the *matK* region was blasted against both NCBI and BOLD databases. DNA barcodes found a 99% sequence homology with the reference *C. sativa* database entries in the NCBI and BOLD databases.

The preliminary basal salt screening compared Murashige and Skoog basal salt medium (MS), Driver and Kuniyuki Walnut basal salt medium (DKW), BABI basal salt medium (BABI) and Gamborg’s B-5 basal salt medium (Gamborg) (Table 1). The explants grown on BABI and Gamborg showed little to no growth after 30 days and had a high mortality rate (Fig. *S3a,b*). Explants grown on DKW had darker and broader leaves, and slightly more growth than the explants grown on MS (Fig. *S3c,d*). Based on these results we hypothesized that DKW could be a more optimal basal salt than MS for growing *Cannabis* in culture.

**Table 1.** Ingredient list for the four basal salt media tested in the preliminary experiment.

| Composition (mg/L) | MS<br>(Murashige and<br>Skoog 1962) | DKW<br>(Driver and<br>Kuniyuki 1984) | BABI<br>(Greenway<br>et al. 2012) | Gamborg<br>(Gamborg et al.<br>1968) |
| --- | --- | --- | --- | --- |
| Ammonium Nitrate | 1650 | 1416 | 320 | 0.0 |
| Ammonium Phosphate | 0.0 | 0.0 | 230 | 0.0 |
| Ammonium Sulfate | 0.0 | 0.0 | 134 | 134 |
| Boric Acid | 6.20 | 4.80 | 3 | 3 |
| Calcium Chloride, Anhydrous | 332.20 | 112.50 | 332.15 | 113.24 |
| Calcium Nitrate | 0.0 | 1367 | 0.0 | 0.0 |
| Cobalt Chloride Hexahydrate | 0.025 | 0.0 | 0.025 | 0.025 |
| Cupric Sulfate Pentahydrate | 0.025 | 0.25 | 0.039 | 0.025 |
| Disodium EDTA Dihydrate | 37.26 | 45.40 | 37.30 | 37.26 |
| Ferrous Sulfate Heptahydrate | 27.80 | 33.80 | 27.80 | 27.80 |
| Magnesium Sulfate, Anhydrous | 180.70 | 361.49 | 122.09 | 122.09 |
| Manganese Sulfate hydrate | 16.90 | 33.50 | 10 | 10 |
| Molybdic Acid (Sodium Salt) Dihydrate | 0.25 | 0.39 | 0.25 | 0.25 |
| Nickel Sulfate Hexahydrate | 0.0 | 0.005 | 0.0 | 0.0 |
| Potassium Iodide | 0.83 | 0.0 | 0.75 | 0.75 |
| Potassium Nitrate | 1900 | 0.0 | 2500 | 2500 |
| Potassium Phosphate, Monobasic | 170 | 265 | 0.0 | 0.0 |
| Potassium Sulfate | 0.0 | 1559 | 0.0 | 0.0 |
| Sodium Phosphate, Monobasic | 0.0 | 0.0 | 150 | 150 |
| Zinc Nitrate Hexahydrate | 0.0 | 17 | 0.0 | 0.0 |
| Zinc Sulfate Heptahydrate | 8.60 | 0.0 | 2 | 2 |
| Total number of components | 14 | 14 | 16 | 14 |
| Total grams per litre | 4.33 | 5.22 | 3.87 | 3.21 |

When comparing explants grown on MS-T05 and DKW-T05, all five cultivars had a larger canopy area on DKW-T05 than on MS-T05, however, these values were only statistically significant for 4/5 of the tested cultivars (Fig. 1*a*; p= < 0.01). The BA-61 cultivar did not have a significant difference in canopy area between the two basal salts (p= 0.21) however a similar trend can be observed (Fig. 1*a*). The average total canopy area of all the cultivars combined for DKW-T05 was significantly greater than MS-T05 at 8.84 cm^2^ and 4.50 cm^2^ respectively (p= < 0.0001). Overall, explants grown on DKW-T05 had a 96.4% increase in canopy area compared to those grown on MS-T05. Qualitative observations found that across all cultivars, explants grown on DKW-T05 had lower incidences of hyperhydricity and leaves that were darker and broader than explants grown on MS-T05 (Fig. 2*a,b*). Leaf morphology on the explants grown on DKW-T05 appeared normal and fully expanded (Fig. *2e*) while those on MS-T05 tended to be long, thin, and curled (Fig. *2f*). Slight yellowing around the edges of explant leaves was observed in both treatments (Fig. *2e,f*).

**Fig. 1.**
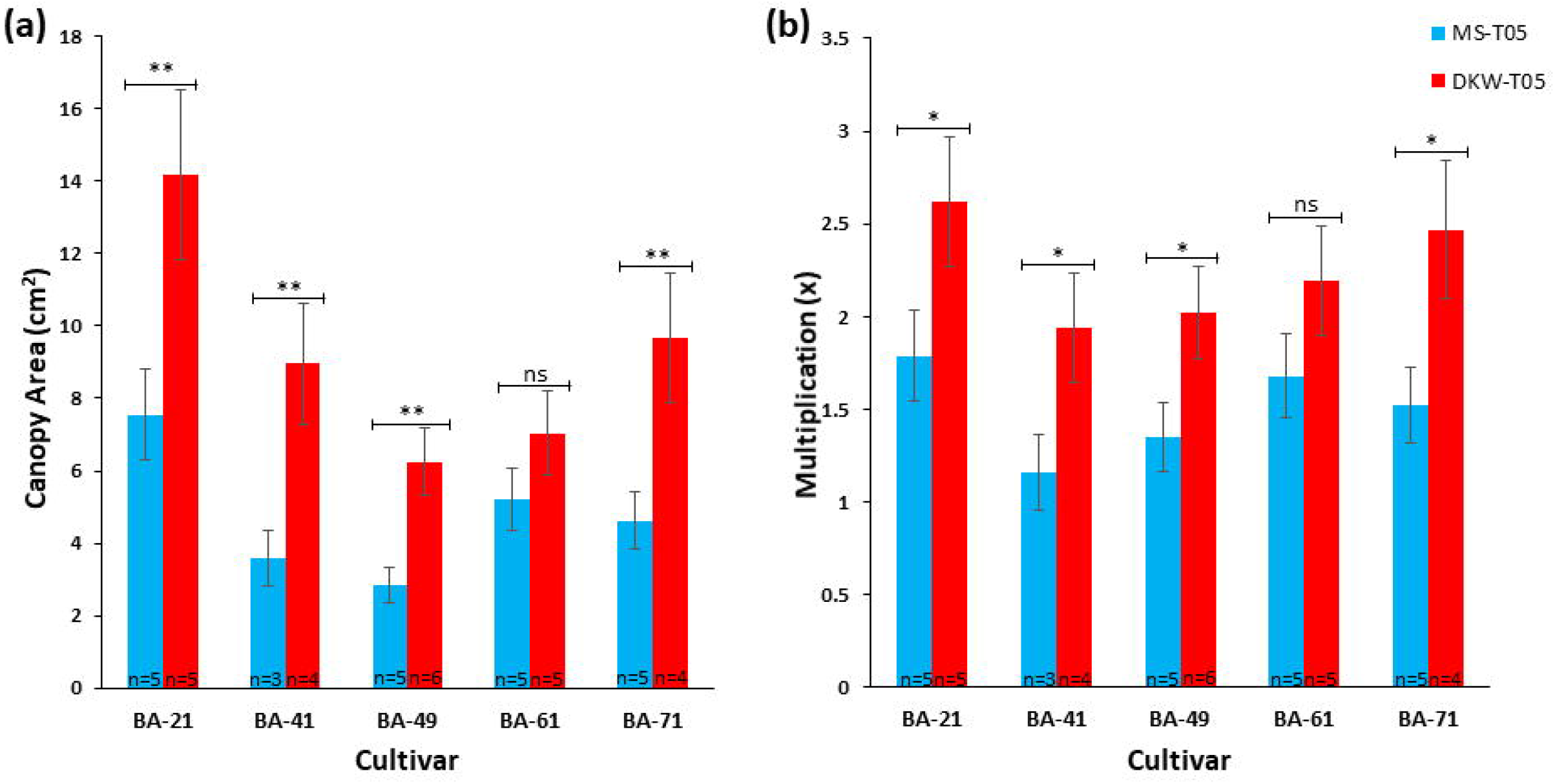
(*a*) Average canopy area (cm^2^) and multiplication rates from three trials conducted using five cultivars (BA-21, BA-41, BA-49, BA-61 and BA-71). (*b*) Average multiplication rate (x) per genotype. * denote significance as determined by two-way ANOVA with a Tukey’s multiple comparison tests (ns p>0.05; * p≤0.05; ** p≤0.01; SAS Studio Software v9.4). Error bars represent standard error of the mean. Explants grown on DKW-T05 generally produced larger canopies and had higher multiplication rates. These results were significant for 80% (4/5) of the cultivars tested.

**Fig. 2.**
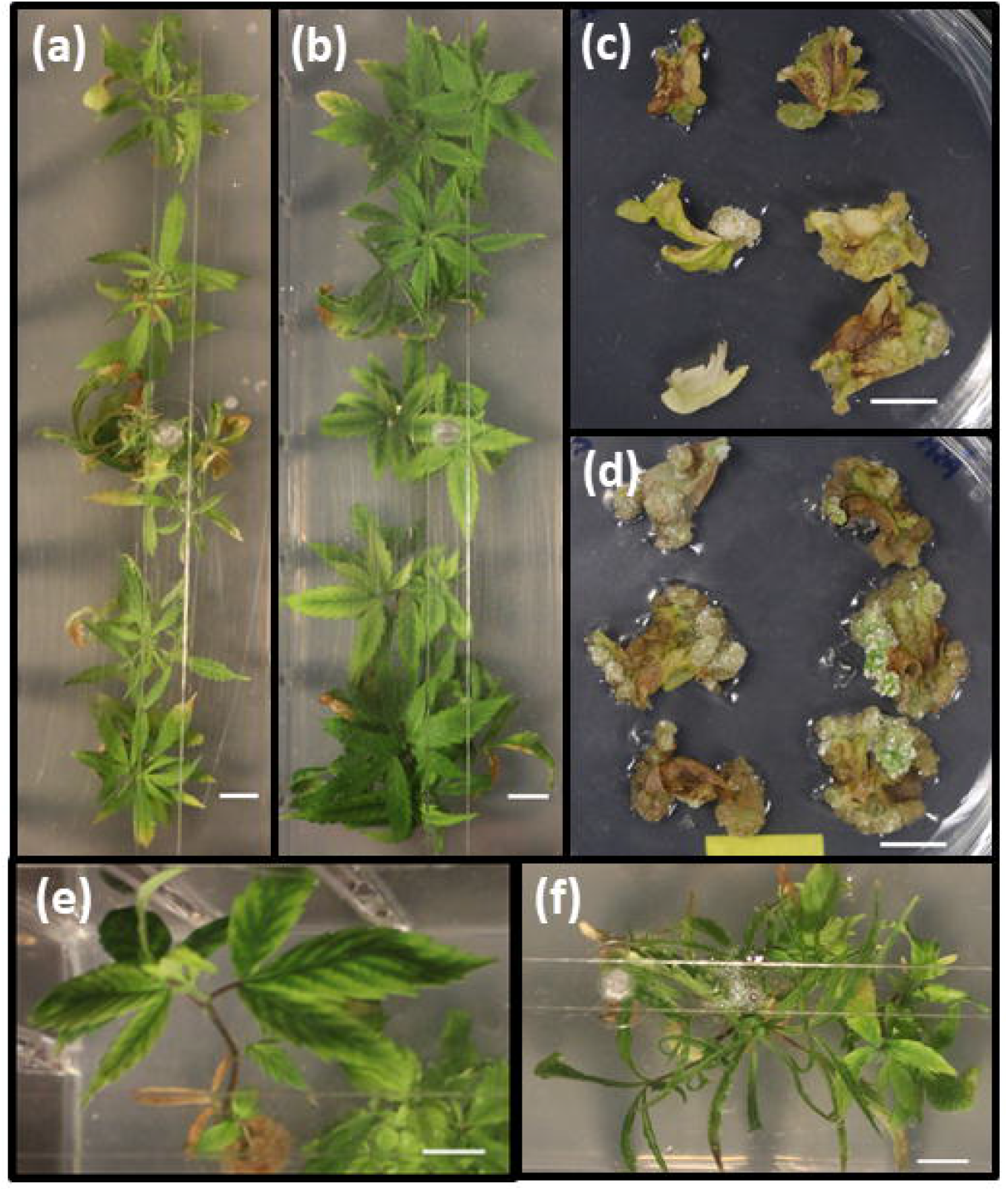
Representative photographs of phenotypic responses in the vegetative growth and callogenesis experiments. (*a*) BA-71 explants grown on MS + 0.5 μM thidiazuron (TDZ). (*b*) BA-71 explants grown on DKW + 0.5 μM TDZ. In 80% of the cultivars tested, explants grown on DKW media had more vigorous growth, less hyperhydricity and broader leaves than those grown on MS media. (*c*) BA-1 leaf disks grown on MS + 10 μM 2,4-D. (*d*) BA-1 leaf disks grown on DKW + 10 μM 2,4-D. Callus growth on DKW media was more vigorous compared to MS media. (*e*) BA-21 explant grown on DKW + 0.5 μM TDZ with yellowing leaf tips. (*f*) BA-21 explant grown on MS + 0.5 μM TDZ with abnormal leaf morphology and yellowing leaf tips. Scale bar: 1 cm.

Similarly, all five cultivars had a greater average multiplication rate on the DKW-T05 medium, but this was only statistically significant for 4/5 of the tested cultivars (Fig. 1*b*; p= < 0.05). The BA-61 cultivar did not have a significant increase in multiplication between the two basal salts (Fig. 1*b*; p= 0.17). The multiplication rate of all the cultivars combined was significantly higher on DKW-T05 than on MS-T05, at 2.23 and 1.48 new 2-node explants per parent explant, respectively (p= <0.0001). On average, the multiplication rate of explants grown on DKW-T05 was 150.7% that of the explants grown on MS-T05.

For the callogenesis experiments no calli were observed on leaf explants cultured in the absence of 2,4 dichlorophenoxyacetic acid (2,4-D) over the experimental period (35 days) on either basal salt. Callus surface area consistently increased over the experimental period in all treatments of 2,4-D. Qualitative observations showed more vigorous and healthier callus growth in 2,4-D treatments cultured on DKW basal salts compared with the respective MS basal salt treatments (Fig. *2c,d*). A comparison of a linear regression of the treatment averages (10, 20 and 30 μM 2,4-D) showed that DKW basal salts resulted in a significantly (p<0.0001) larger callus surface area than growth on MS basal salts (Fig. 3*a*). Similar trends were observed for both trials; however, the magnitude of the callus surface area was consistently lower in the second trial. Leaf explants cultured on MS and DKW basal salts in the absence of 2,4-D (control) did not show callus formation. Control tissues began necrosis after 4 weeks and were dead by the end of the experiment after 5 weeks. Callus formation was observed at all three treatment levels (10, 20 and 30 μM 2,4-D) on both basal salts tested. An analysis of variance (ANOVA) with a post-hoc student’s t-test found that at all three concentrations of 2,4-D, callus mass was significantly greater (p< 0.05) on DKW basal salts compared to the same treatment on MS. This trend was observed in both trials. In the first trial, 10 μM concentration of 2,4-D showed the greatest callus production of all explants grown on DKW basal salts (p<0.05; Fig. 3*b*), averaging 358 mg per explant. There was, however, no significant concentration dependence for callus production observed within DKW treatments in the second trial. The absolute value obtained for callus masses were greater in the first trial compared to the second trial, as was observed in the callus surface area experiments.

**Fig. 3.**
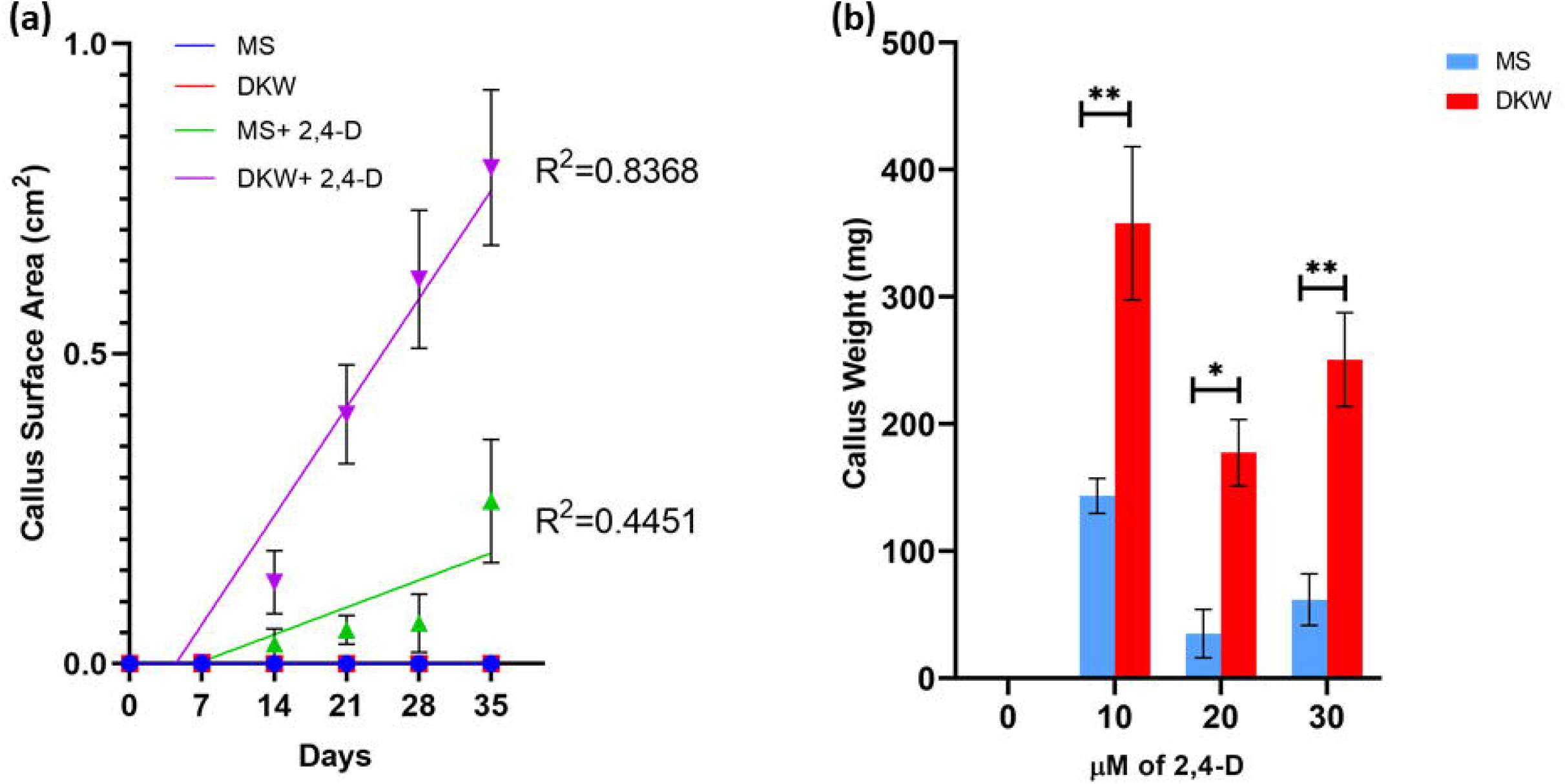
Trial 1 results for surface area and callus weight of BA-1 leaf disks cultured on MS and DKW media with 2,4 dichlorophenoxyacetic acid (2,4-D). Responses in trial 1 and trial 2 showed similar trends, the overall magnitude of response was lower in trial 2. (*a*) Linear regression of callus surface area. MS and DKW represent controls with no 2,4-D and did not produce any callus. MS + 2,4-D and DKW + 2,4-D represent the average surface area responses observed across all 2,4-D treatment levels (10, 20, and 30 μM). Comparison of linear regressions found that the callus response to DKW + 2,4-D treatments was significantly greater than the MS + 2,4-D average response (p<0.001). (*b*) Callus weight in milligrams (mg) per leaf disk. * denote significance as determined by a two-way ANOVA with a Tukey’s multiple comparison tests (ns p>0.05; * p≤0.05; ** p≤0.01; Prism v8.3). Similar trends were observed for the second trial. Error bars represent standard error of the mean.

## Discussion

Previous studies have reported that MS medium supplemented with TDZ is suitable for micropropagation of drug-type *Cannabis* (Chandra et al. 2009; Lata et al. 2009b; Wróbel et al. 2020), with the highest multiplication rate reported to be 12.6 using MS + 0.5 μM TDZ (Lata et al. 2009a). In the present study, nodal explants cultured on this concentration of TDZ resulted in a much lower multiplication rate of 1.48 averaged across five cultivars. One possible explanation for this difference is that there is strong genotypic variation in the response of different cultivars, a phenomenon that has previously been demonstrated in industrial hemp (Slusarkiewicz-Jarzina et al. 2005; Chaohua et al. 2016). The analysis of 11 different SSR regions within the 6 cultivars revealed considerable diversity in the experimental population (Fig. S1). Strong phenotypic variations have not only been seen in industrial hemp, but also in drug-type *Cannabis* and have fueled the ongoing debate surrounding the *indica* and *sativa* nomenclature of the genus *Cannabis* (reviewed by McPartland (2018)). While The National Institute on Drug Abuse (NIDA) remains the main supplier for research-grade *Cannabis* in the United States and a recent report suggested that NIDA- supplied *Cannabis* is genetically more similar to industrial hemp than to many drug-type *Cannabis* cultivars on the market (Schwabe et al. 2019). This difference that is echoed by findings that dispensary-sold drug-type *Cannabis* and *Cannabis* obtained for research purposes from the NIDA differ in their chemical compositions (Vergara et al. 2017). The use of a single cultivar such as MX used by Lata et al. (2009a) may result in findings that are highly specific to an individual cultivar and not necessarily applicable to all drug-type cultivars, such as those used in this current study which were not supplied by the NIDA. Given that there are multiple reports of plant regeneration in industrial hemp (Slusarkiewicz-Jarzina et al. 2005; Plawuszewski et al. 2006; Chaohua et al. 2016; Smýkalová et al. 2019), a possible explanation for the discrepancy between our findings and some existing publications is that these may share closer genetic relatedness to industrial hemp making them more amenable to micropropagation.

While genetic differences could explain some discrepancies between our study and the findings of Lata et al. (2009a), it is not the only major difference. Study design plays an important role in the outcome of explant multiplication. Micropropagation is generally divided into five stages including Stage 0: Selection/maintenance of parent plant material, Stage 1: Initiation of cultures, Stage 2: Multiplication of shoots, Stage 3: Shoot elongation and rooting, and Stage 4: Acclimatization (Murashige 1974; George et al. 2008) (Fig. S4). In most *Cannabis* micropropagation studies (Lata et al. 2009a; Smýkalová et al. 2019), nodal explants from plants growing in a greenhouse or growth room are initiated into culture and various media are screened to identify the optimal medium for shoot multiplication. These shoots are then transferred to rooting medium to induce root formation and transferred back to the growth facilities. As such, these studies have been designed to optimize the multiplication rate during the initiation phase (stage 1) rather than the multiplication phase (stage 2). The present study was designed to work with and improve the micropropagation of stage 2 explants. In other species, it has been well established that the optimal conditions required during stage 1 can be different from the optimal conditions for stage 2. Furthermore, it is common to see a flush of growth during the initiation phase, followed by slower and more sporadic growth until cultures stabilize (Murashige 1974; Vardja and Vardja 2001). The growth differences observed between these two stages may explain why we observed a progressive decline and frequent explant death when maintaining cultures on a medium that was optimized for stage 1 micropropagation. Tissue culture’s main benefit is the possibility to rapidly multiply plants through repeated cycles of subculturing during stage 2, while multiplication protocols using stage 1 growth followed by immediate rooting provides only a portion of the potential benefits. As such, the optimization of stage 2 micropropagation is critical for commercial application.

The slow decline of plants when cultured on MS suggests that something in the medium is negatively affecting the plants and the stress is accumulating over time. While MS medium is widely used and very effective for many plant species, there are others that do not do well on this specific nutrient mixture (Gamborg et al. 1968; Lloyd and McCown 1980; Driver and Kuniyuki 1984; Greenway et al. 2012) This has led to the development of many different basal media specifically developed for specific plants. In the current study, four different basal salts were evaluated in a preliminary screen: MS (Murashige and Skoog 1962), DKW (Driver and Kuniyuki 1984), Gamborg (Gamborg et al. 1968), and BABI (Greenway et al. 2012). Of these four media, MS and DKW produced the best results, while Gamborg and BABI resulted in very poor growth and were not further evaluated. The composition of these media is summarized in detail by Phillips and Garda (2019) but some of the major differences include the amount and relative proportions of various salts and more specifically the amount and type of nitrogen. It is important to note that of these four, MS and DKW contain the highest amount of total salts, with 4.3 g/L and 5.32 g/L (Table 1) respectively, suggesting that *Cannabis* prefers a nutrient-rich medium.

Between DKW and MS basal medium, DKW was superior and resulted in higher multiplication rates, larger canopy area, and overall healthier looking plants (Fig. 2*a,b*). While plants on MS medium were very hyperhydric and had extremely deformed leaves that often failed to expand, plants cultured on DKW were visibly less hyperhydric with normally developed leaves. However, it is important to note that the symptoms observed in the present study were based on the first subculture from plants maintained on DKW medium. From previous experience, the symptoms were more severe when plants were maintained on MS for multiple subcultures and this often resulted in plant death. For example, while the BA-61 cultivar did not show any significant difference when grown on DKW-T05 compared to MS-T05, we previously found that maintaining BA-61 on either MS or MS-T05 over multiple subcultures resulted in a decline in explant health and almost killed our entire BA-61 germplasm. When these plants were transferred to DKW based media, they recovered and have been subcultured many times without a significant decline. A similar problem with walnut led to the development of DKW: while the researchers were initially able to obtain multiple shoot formation from walnut on MS and other basal media, multiple successive subcultures led to callus rather than shoot formation. Within ten subcultures they were no longer able to produce viable shoots (Driver and Kuniyuki 1984). This is very similar to observations with *Cannabis*, where MS-T05 facilitated explant growth during initial transfers, but over time we observed increased callusing, poor shoot quality, and in some cases explant death.

In comparison to MS basal salts, DKW has similar ammonium to nitrate ratio but has less total nitrogen (Driver and Kuniyuki 1984; Phillips and Garda 2019). While this may contribute to the difference in plant growth, DKW is also higher in total salt levels and has a significantly different composition. Some major differences to note are that DKW includes much higher levels of sulphur (~7x), calcium (~3x), and Copper (10x) (Table 2). Further research is needed to determine which factors are responsible for the improved growth in *Cannabis* and how this can be further improved. While DKW was superior to MS, some explants had yellowing around the edge of their leaves on both media, suggesting that there is still a nutrient imbalance and further optimization may be beneficial. Once the basal medium is fully optimized, it would also be useful to re-examine the type and levels of plant growth regulators in order to create a medium that is optimized for the majority of the *Cannabis*’ time in culture.

**Table 2.** The amounts of each element found in MS and DKW basal salt media. The total amounts of nitrate, ammonium, sulfate, and phosphate are also listed.

| Composition (mg/L) | MS<br>(Murashige and<br>Skoog 1962) | DKW<br>(Driver and Kuniyuki<br>1984) |
| --- | --- | --- |
| C | 12.02 | 14.65 |
| H | 90.03 | 80.72 |
| O | 2119.25 | 2610.04 |
| N | 843.82 | 734.24 |
| P | 38.69 | 60.31 |
| K | 783.77 | 775.65 |
| Ca | 119.96 | 374.58 |
| Mg | 36.48 | 72.98 |
| S | 55.52 | 393.44 |
| Zn | 1.96 | 3.74 |
| Mo | 0.099 | 0.16 |
| B | 1.08 | 0.84 |
| Cu | 0.0064 | 0.064 |
| Mn | 5.49 | 10.87 |
| Na | 4.65 | 5.68 |
| Fe | 5.59 | 6.79 |
| Ni | 0.0 | 0.0012 |
| Cl | 212.25 | 71.88 |
| Co | 0.0062 | 0.0 |
| I | 0.63 | 0.0 |
| NH<sub>4</sub> | 372.08 | 319.31 |
| NO<sub>3</sub> | 2443.72 | 2137.24 |
| SO<sub>4</sub> | 166.31 | 1178.81 |
| PO<sub>4</sub> | 118.63 | 184.92 |

Numerous research groups have also explored the formation and subsequent development of callus in *Cannabis*, mostly with the goal of plant regeneration (Raharjo et al. 2006; Chandra et al. 2009; Lata et al. 2010; Farag 2014; Movahedi et al. 2015; Chaohua et al. 2016; Piunno et al. 2019). These studies have largely used MS basal salts and to the authors’ knowledge, none have explored the use of DKW to produce callus (Richez-Dumanois et al. 1986; Mandolino and Ranalli 1999; Slusarkiewicz-Jarzina et al. 2005; Raharjo et al. 2006; Chandra et al. 2009; Lata et al. 2009a, 2010; Movahedi et al. 2015; Chaohua et al. 2016; Smýkalová et al. 2019; Piunno et al. 2019). Given that whole plants did not perform well on MS and preferred DKW, these two basal media were evaluated for callogenesis. Similar to whole shoot growth, the surface area of 2,4-D induced callus increased significantly faster on DKW compared to MS over the period of study (35 days; Fig. 3*a*). These findings are further supported by the significantly larger callus mass found at all 2,4-D treatment levels on DKW at the end of the 35-day study period (Fig. 3*b*). This suggests that 2,4-D mediated callogenesis is a product of the interaction between basal salts and PGRs, a link that has not previously been explored in *Cannabis*. The results of this study suggest that for the development of callus using 2,4-D, DKW basal salts are preferable to MS basal salts. However, given that no regeneration was observed, it is not known if DKW would be beneficial for this application. While most research groups have used MS in the proliferation of callus cultures, DKW and other media should not be overlooked as a potentially better option for callogenesis and plant regeneration. Further work is warranted to determine whether the auxin-DKW interaction leads to greater callus proliferation in all commonly used auxins, or if this enhanced callogenesis on DKW occurs in an auxin-specific manner, as shown by the response to 2,4-D in this experiment.

Based on our research, the previously reported *Cannabis* growth medium of MS + 0.5 μM TDZ is not optimal for all cultivars in stage 2 micropropagation. *Cannabis* grown on DKW-T05 displayed fewer physiological abnormalities, better multiplication rates, larger canopy area and broader leaves in four out of five genotypes evaluated, in addition to improved callogenesis. While DKW-T05 is an improvement over MS-T05, the multiplication rate remains relatively low. Further improvements in media composition are likely possible and DKW basal salt medium provides a good starting point for this objective. Continued research will ultimately help to make micropropagation a more effective approach for germplasm maintenance and clean plant production in *Cannabis sativa*.

## Acknowledgements

The authors gratefully acknowledge our industry partners, Canopy Growth Corporation and Hexo Corp. for the use of their plant material. The financial support of the Natural Sciences and Engineering Research Council of Canada (Grant No. RGPIN-2016-06252) is also gratefully acknowledged. This manuscript has been released as a pre-print at bioRxiv (Page et al. 2020).

## Abbreviations

2,4-D: 2,4 dichlorophenoxyacetic acid
ANOVA: analysis of variance
BABI: BDS as modified at Arkansas Bioscience Institute
CBD: cannabidiol
CCDB: Canadian Centre for DNA Barcoding
DKW: Driver and Kuniyuki Walnut
MS: Murashige and Skoog
NIDA: National Institute on Drug Abuse
PAR: photosynthetically active radiation
PGR: plant growth regulator
PPM: Plant Preservative Mixture
SSR: simple sequence repeats
TDZ: thidiazuron
THC: Δ^9^-tetrahydrocannabinol

---

**Supplemental Fig. S1.**
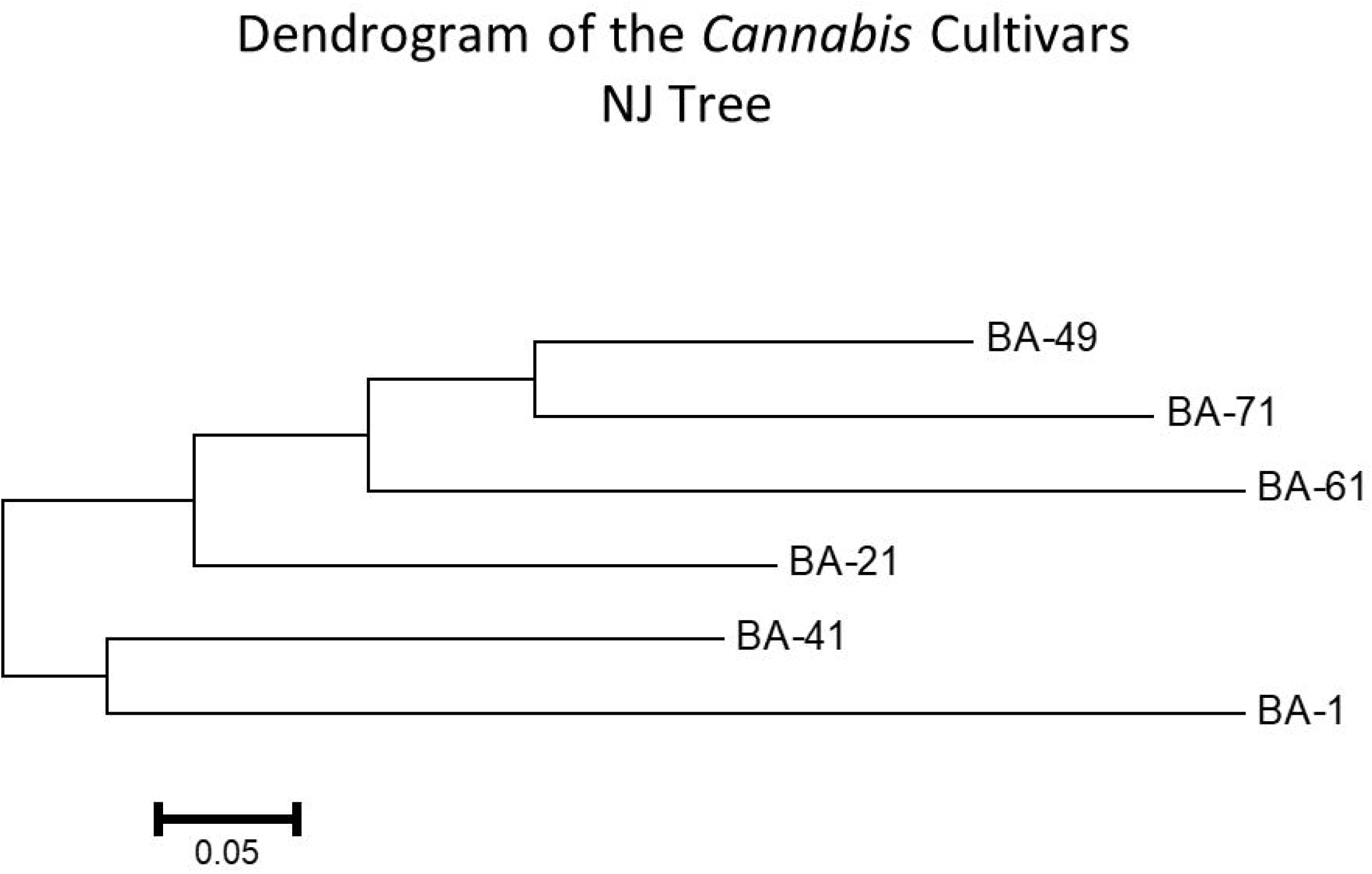
Dendrogram of the *Cannabis* cultivars used in the vegetative and callogenesis studies show clustering of two genetically distinct groups within the species. Dendrogram was generated from SSR sequence data using the methods as first reported by (Alghanim and Almirall 2003). DNA barcodes of BA-1, BA-21, BA-41, BA-49, BA-61 and BA-71 were obtained from the *MatK* gene region with a 99% sequence agreement with reference *Cannabis sativa* database entries. BA-1 was exclusively used for callogenesis experiments while BA-21, BA-41, BA-49, BA-61 and BA-71 were used exclusively in the vegetative growth studies.

**Supplemental Fig. S2.**
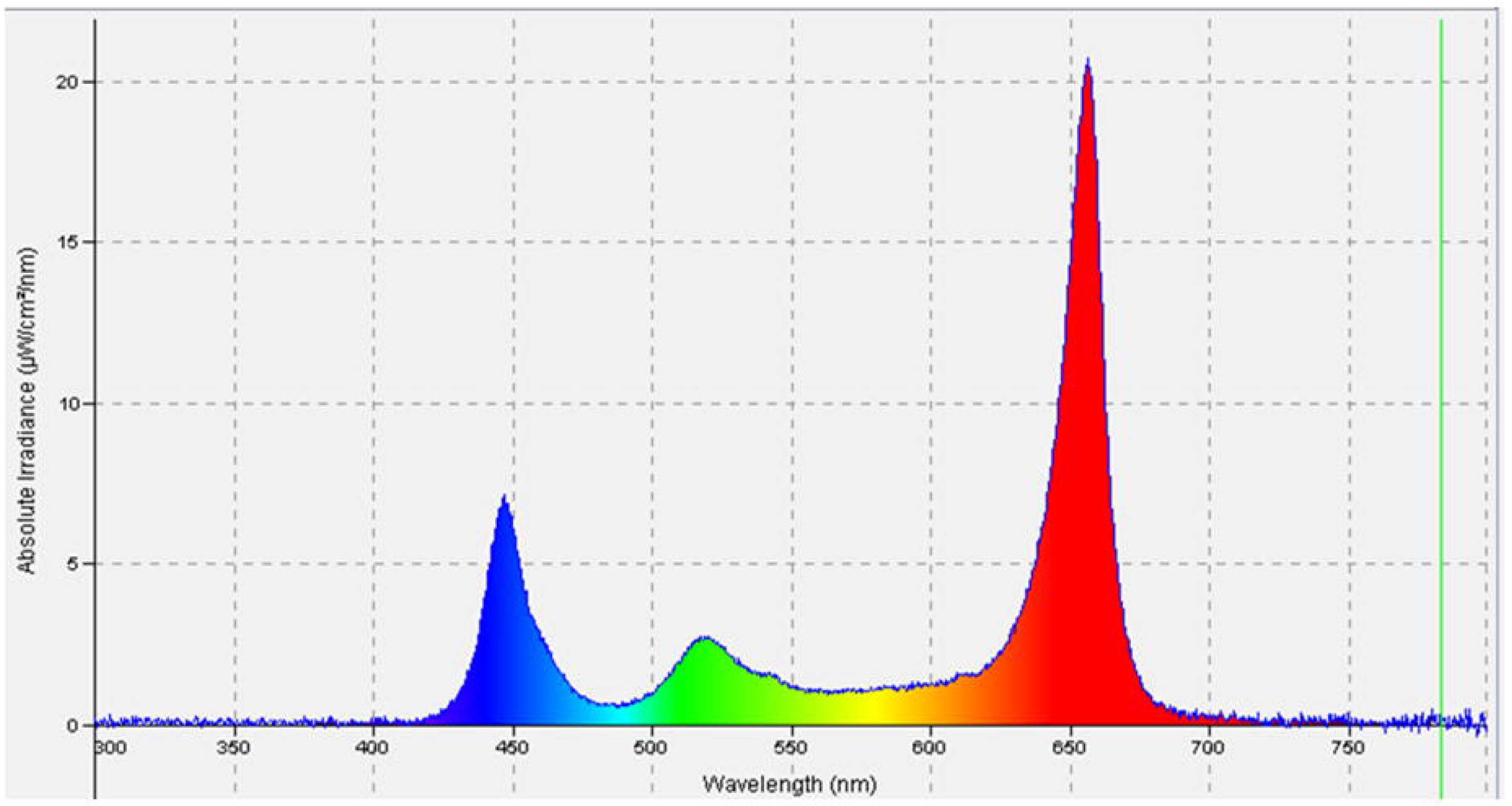
A representative light spectrum of the lighting used in the controlled environment growth chamber. The average photosynthetically active radiation (PAR) over the experimental area was 41 ± 4 μmol s^−1^ m^−2^ using an OceanOptics. Average PAR was calculated using Excel ™.

**Supplemental Fig. S3.**
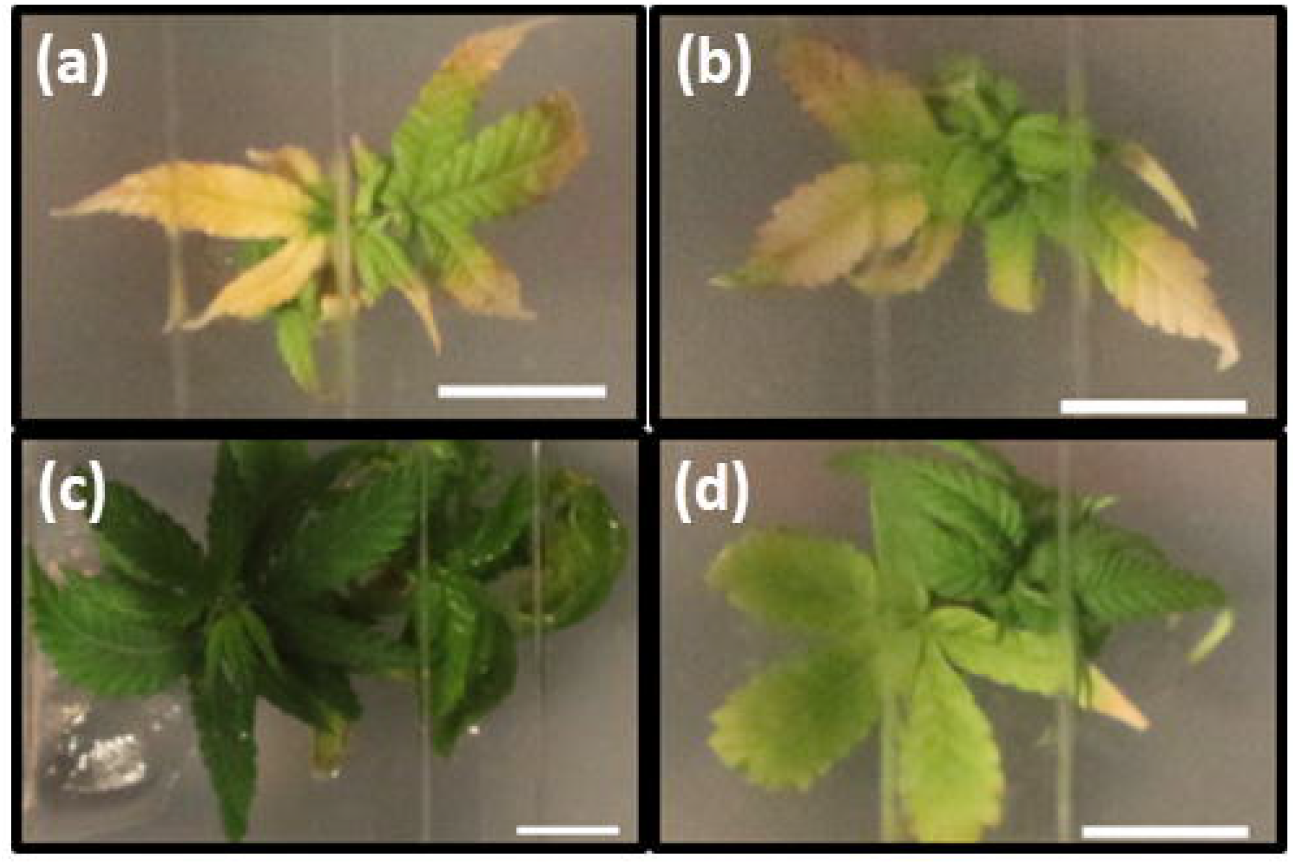
Representative photographs of the phenotypic response to different basal salt media. (*a*) BA-1 explant on Gamborg basal salt medium. (*b*) BA-1 explant on BABI basal salt medium. (*c*) BA-1 explant on DKW basal salt medium. (*d*) BA-1 explant on MS basal salt medium. Scale bar: 1 cm.

**Supplemental Fig. S4.**
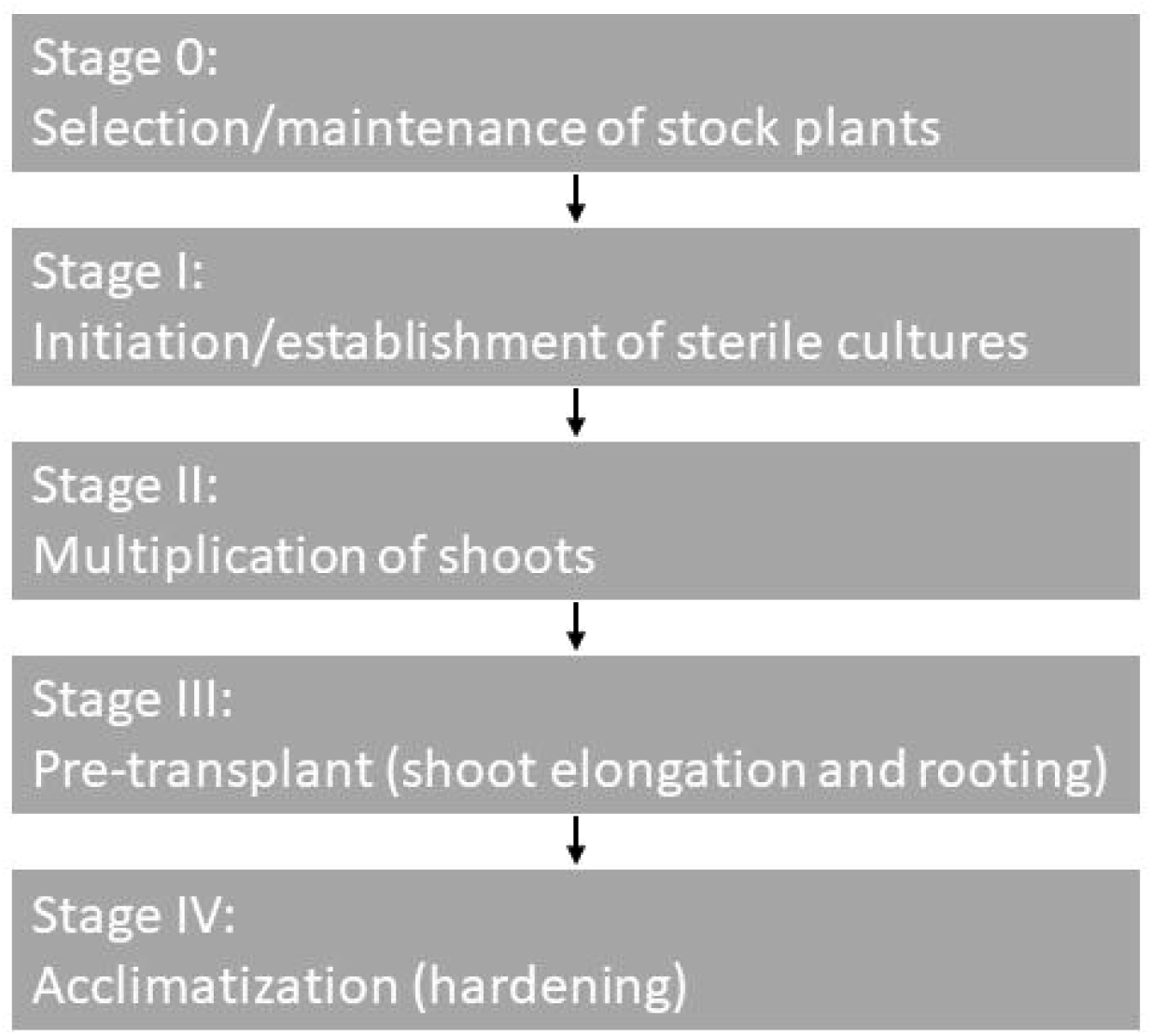
Diagram of the stages of micropropagation.

## References

Alghanim, H.J., and Almirall, J.R. 2003. Development of microsatellite markers in Cannabis sativa for DNA typing and genetic relatedness analyses. Anal. Bioanal. Chem. 376(8): 1225–1233. doi:10.1007/s00216-003-1984-0.

Chandra, S., Lata, H., Khan, I.A., and ElSohly, M.A. 2011. Photosynthetic response of Cannabis sativa L., an important medicinal plant, to elevated levels of CO2. Physiol. Mol. Biol. Plants 17(3): 291–295. doi:10.1007/s12298-011-0066-6.

Chandra, S., Lata, H., Khan, I.A., and ElSohly, M.A. 2017. Chapter 3: Cannabis sativa L.: Botany and Horticulture. In Cannabis sativa L. – Botany and Biotechnology. Edited by S. Chandra, H. Lata, and M.A. ElSohly. Springer International Publishing, Cham. doi:10.1007/978-3-319-54564-6.

Chandra, S., Lata, H., Mehmedic, Z., Khan, I.A., and ElSohly, M.A. 2009. Assessment of Cannabinoids Content in Micropropagated Plants of Cannabis sativa L. and their Comparison with Vegetatively Propagated Plants and Mother Plant at Different Stages of Growth. Planta Med. 75(04): 743–750. doi:10.1055/s-2009-1216442.

Chaohua, C., Gonggu, Z., Lining, Z., Chunsheng, G., Qing, T., Jianhua, C., Xinbo, G., Dingxiang, P., and Jianguang, S. 2016. A rapid shoot regeneration protocol from the cotyledons of hemp (Cannabis sativa L.). Ind. Crops Prod. 83: 61–65. Elsevier B.V. doi:10.1016/j.indcrop.2015.12.035.

Clarke, R.C., and Merlin, M.D. 2013. Classical and Molecular Taxonomy. In Cannabis: Evolution and Ethanobotany. University of California Press, Berkeley. pp. 311–331.

Driver, J., and Kuniyuki, A. 1984. In vitro propagation of Paradox walnut rootstock. HortScience 19(4): 507–509.

Farag, S. 2014. Cannabinoids production in Cannabis sativa L.: An in vitro approach [Dissertation]. Technischen Universität Dortmund.

Gamborg, O.L., Miller, R.A., and Ojima, K. 1968. Nutrient requirements of suspension cultures of soybean root cells. Exp. Cell Res. 50: 150–158.

George, E.F., Hall, M.A., Klerk, G.J. De, and Debergh, P.C. 2008. Micropropagation: Uses and methods. In Plant Propagation by Tissue Culture 3rd Edition. Springer Netherlands. pp. 29–64. doi:10.1007/978-1-4020-5005-3_2.

Government of Canada. 2015. Industrial Hemp Regulations Règlement sur le chanvre industriel. Ministry of Justice, Ottawa.

Greenway, M.B., Phillips, I.C., Lloyd, M.N., Hubstenberger, J.F., and Phillips, G.C. 2012. A nutrient medium for diverse applications and tissue growth of plant species in vitro. Vitr. Cell. Dev. Biol. – Plant 48: 403–410. doi:10.1007/s11627-012-9452-1.

Hillig, K.W. 2004. A chemotaxonomic analysis of terpenoid variation in Cannabis. Biochem. Syst. Ecol. 32(10): 875–891. doi:10.1016/j.bse.2004.04.004.

Hillig, K.W. 2005. Genetic evidence for speciation in Cannabis (Cannabaceae). Genet. Resour. Crop Evol. 52(2): 161–180. doi:10.1007/s10722-003-4452-y.

Ivanova, N. V, and Grainger, C. 2007. Dye terminator sequencing for COI for the 3730xl DNA Analyzer. Guelph. Available from http://ccdb.ca/site/wp-content/uploads/2016/09/CCDB_Sequencing.pdf.

Kuzmina, M., and Ivanova, N. 2011a. PCR Amplification for Plants and Fungi. Guelph. Available from http://ccdb.ca/site/wp-content/uploads/2016/09/CCDB_Amplification-Plants.pdf.

Kuzmina, M., and Ivanova, N. 2011b. Primer Sets for Plants and Fungi. Guelph. Available from http://ccdb.ca/site/wp-content/uploads/2016/09/CCDB_PrimerSets-Plants.pdf.

Kuzmina, M., Ivanova, N., and Fazekas, A. 2011. CCDB Protocols, Glass Fiber Plate DNA Extraction Protocol for Plants, Fungi, Echinoderms and Mollusks. Guelph. Available from http://ccdb.ca/site/wp-content/uploads/2016/09/CCDB_DNA_Extraction-Plants.pdf.

Lata, H., Chandra, S., Khan, I., and ElSohly, M.A. 2009a. Thidiazuron-induced high-frequency direct shoot organogenesis of Cannabis sativa L. Vitr. Cell. Dev. Biol. – Plant 45(1): 12–19. doi:10.1007/s11627-008-9167-5.

Lata, H., Chandra, S., Khan, I.A., and Elsohly, M.A. 2009b. Propagation through alginate encapsulation of axillary buds of Cannabis sativa L. – An important medicinal plant. Physiol. Mol. Biol. Plants 15(1): 79–86. doi:10.1007/s12298-009-0008-8.

Lata, H., Chandra, S., Khan, I.A., and ElSohly, M.A. 2010. High Frequency Plant Regeneration from Leaf Derived Callus of High Δ 9 – Tetrahydrocannabinol Yielding Cannabis sativa L. Planta Med. 76(14): 1629–1633. doi:10.1055/s-0030-1249773.

Lata, H., Chandra, S., Techen, N., Khan, I.A., and ElSohly, M.A. 2016. In vitro mass propagation of Cannabis sativa L.: A protocol refinement using novel aromatic cytokinin meta-topolin and the assessment of eco-physiological, biochemical and genetic fidelity of micropropagated plants. J. Appl. Res. Med. Aromat. Plants 3(1): 18–26. Elsevier GmbH. doi:10.1016/j.jarmap.2015.12.001.

Lloyd, G., and McCown, B. 1981. Commercially-feasible micropropagation of Mountain Laurel, Kalmia latifolia, by use of shoot tip culture. Proc. Int. Plant Propagators’ Soc. 30: 421–427.

Mandolino, G., and Ranalli, P. 1999. Advances in biotechnological approaches for hemp breeding and industry. In Advances in hemp research. pp. 185–212.

McPartland, J.M. 2018. Cannabis Systematics at the Levels of Family, Genus, and Species. Cannabis Cannabinoid Res. 3(1): 203–212. doi:10.1089/can.2018.0039.

Movahedi, M., Ghasemi-Omran, V.-O., and Torabi, S. 2015. The effect of different concentrations of TDZ and BA on in vitro regeneration of Iranian cannabis (Cannabis sativa) using cotyledon and epicotyl explants. J. Plant Mol. Breed. 3(2): 20–27. doi:10.22058/jpmb.2015.15371.

Murashige, T. 1974. Plant Propagation Through Tissue Cultures. Annu. Rev. Plant Physiol. 25(1): 135–166. doi:10.1146/annurev.pp.25.060174.001031.

Murashige, T., and Skoog, F. 1962. A Revised Medium for Rapid Growth and Bio Assays with Tobacco Tissue Cultures. Physiol. Plant. 15(3): 473–497. doi:10.1111/j.1399-3054.1962.tb08052.x.

Page, S.R.G., Monthony, A.S., and Jones, A.M.P. 2020. Basal media optimization for the micropropagation and callogenesis of Cannabis sativa L. bioRxiv [Preprint]: 1–23. doi:10.1101/2020.02.07.939181.

Phillips, G., and Garda, M. 2019. Plant tissue culture media and practices: an overview. Vitr. Cell. Dev. Biol. – Plant 55(3): 242–257. doi:10.1007/s11627-019-09983-5.

Piunno, K., Golenia, G., Boudko, E.A., Downey, C., and Jones, A.M.P. 2019. Regeneration of shoots from immature and mature inflorescences of Cannabis sativa. Can. J. Plant Sci. 99(4): 556–559. doi:10.1139/cjps-2018-0308.

Plawuszewski, M., Lassociński, W., and Wielgus, K. 2006. Regeneration of Polish cultivars of monoecious hemp (Cannabis sativa L.) grown in vitro. In Renewable Resources and Plant Biotechnology, November 5. pp. 149–154.

Raharjo, T.J., Eucharia, O., Chang, W.-T., and Verpoorte, R. 2006. Callus Induction and Phytochemical Characterization of Cannabis sativa Cell Suspension Cultures. Indones. J. Chem. 6(1): 70–74. doi:10.22146/ijc.21776.

Richez-Dumanois, C., Braut-Boucher, F., Cosson, L., and Paris, M. 1986. Multiplication végétative in vitro du chanvre (Cannabis sativa L.). Application à la conservation des clones sélectionnés. Agronomie 6(5): 487–495. doi:10.1051/agro:19860510.

Schuelke, M. 2000. An economic method for the fluorescent labeling of PCR fragments. Nat. Biotechnol. 18(2): 233–234. doi:10.1038/72708.

Schwabe, A.L., Hansen, C.J., Hyslop, R.M., and McGlaughlin, M.E. 2019. Research grade marijuana supplied by the National Institute on Drug Abuse is genetically divergent from commercially available Cannabis. bioRxiv [Preprint] 7(3): 31–38. doi:10.1101/592725.

Slusarkiewicz-Jarzina, A., Ponitka, A., and Kaczmarek, Z. 2005. Influence of cultivar, explant source and plant growth regulator on callus induction and plant regeneration of Cannabis sativa L. Acta Biol. Cracoviensia Ser. Bot. 47(2): 145–151.

Small, E. 1975. Morphological variation of achenes of Cannabis. Can. J. Bot. 53(10): 978–987. doi:10.1139/b75-117.

Small, E., and Cronquist, A. 1976. A Practical and Natural Taxonomy for Cannabis. Taxon 25(4): 405–435.

Smýkalová, I., Vrbová, M., Cvečková, M., Plačková, L., Žukauskaitė, A., Zatloukal, M., Hrdlicka, J., Plíhalová, L., Doležal, K., and Griga, M. 2019. The effects of novel synthetic cytokinin derivatives and endogenous cytokinins on the in vitro growth responses of hemp (Cannabis sativa L.) explants. Plant Cell, Tissue Organ Cult. 139(2): 381–394. Springer Netherlands. doi:10.1007/s11240-019-01693-5.

Vardja, R., and Vardja, T. 2001. The effect of cytokinin type and concentration and the number of subcultures on the multiplication rate of some decorative plants. Proc. Est. Acad. Sci. Biol. 50(1): 22–31.

Vergara, D., Bidwell, L.C., Gaudino, R., Torres, A., Du, G., Ruthenburg, T.C., Decesare, K., Land, D.P., Hutchison, K.E., and Kane, N.C. 2017. Compromised External Validity: Federally Produced Cannabis Does Not Reflect Legal Markets. Sci. Rep. 7(July 2016): 1–8. Nature Publishing Group. doi:10.1038/srep46528.

Wróbel, T., Dreger, M., Wielgus, K., and Słomski, R. 2020. Modified Nodal Cuttings and Shoot Tips Protocol for Rapid Regeneration of Cannabis sativa L. J. Nat. Fibers: 1–10. Taylor & Francis. doi:10.1080/15440478.2020.1748160.

